# ImageJ Macro for Counting Stationary vs. Moving Items in Videos

**DOI:** 10.1101/292417

**Authors:** T. Douglas, R. Davalos

## Abstract

In microfluidic devices, it is often necessary to determine whether cells or particles are stationary or moving. Here we outline the development of an ImageJ macro that can be used to analyze a video of moving, fluorescently labeled particles or cells, and determine which objects are stationary and which objects are moving at each point in time, providing information on the percentage of cells moving as well as their median radius.

## INTRODUCTION

ImageJ is an imaging software designed by the National Institutes of Health for the purpose of providing an open platform for image analysis. This software includes many built in functions that researchers can utilize, including image formatting, cell counting, as well as documentation for built-in methods that can be accessed using simple Java programming. (1) (2). ImageJ also provides a method to record macros, which makes it easier for individual researchers to develop their own code. In addition, there is an online repository for these methods where other researchers can use the same code for their own work.(3) A common difficulty that arises in analyzing microfluidic data is the problem of determining whether or not objects in a video are moving. This is particularly important in applications of dielectrophoresis, where cells are often trapped or immobilized in part of device, and quantitatively determining the fraction of cells immobilized as a function of applied frequency and voltage becomes critical. Other applications, such as microfluidic chemotaxis assays, devices for fluid dynamics studies using fluorescent beads as flow markers, and CTC sorting applications could also benefit from this software.(4)(5) In our research, we are developing a microfluidic device that can be used for separating highly similar cell types. In our device, we have an interest in determining in what percentage of the cells are trapping at each point in time, correlating with a specific voltage and frequency. In particular, we are interested in understanding if there is a difference in trapping efficiency between two cell types, indicating dielectrophoretic selectivity (6)(7)(8). The current method of analyzing dielectrophoresis separation results is to count the number of cells in each separated batch after the experiment. This is a time consuming process.

The macro presented here provides a method of taking videos obtained during microfluidic dielectrophoresis experiments and extracting in each frame of the video which color cells are moving vs. stationary, how big the moving vs. stationary cells are, and what percentage of each cell type is trapped. An example is presented from our current research. We predict that this application could have value for groups trying to quantify cellular behavior in microdevices.

## MATERIALS AND METHODS

### ImageJ Commands

As with all ImageJ macros, this file should be saved as a text file and opened through the ImageJ program. “Ctrl-R” initializes the program once open, and the image stack of interest should be saved in a folder by itself.

The code is designed to work on an image stack of fluorescent cell or bead images, either single channel or multichannel.

The basic parameters for the code development were developed using the macro record function. “Plugins>Macros>Record”.

From this, a basic outline of the code was developed, following existing ImageJ protocols.(9)(10)

The macro uses several packages from ImageJ including the Dialog package for providing a window for user commands.First, the command “getString(text question, text answer)” is used. This is a command that opens a quick dialog box with a single line of question and input without having to use the full Dialog command. Then, the image sequence of interest is opened. The commands File.getParent(source) and File.getName(source) are used to split the full path into the file name and the path without the filename. This is used in multiple places in the code to query the user. The code can be defined as functioning in eight distinct steps, as shown in Figure 1.

**Figure 1:**
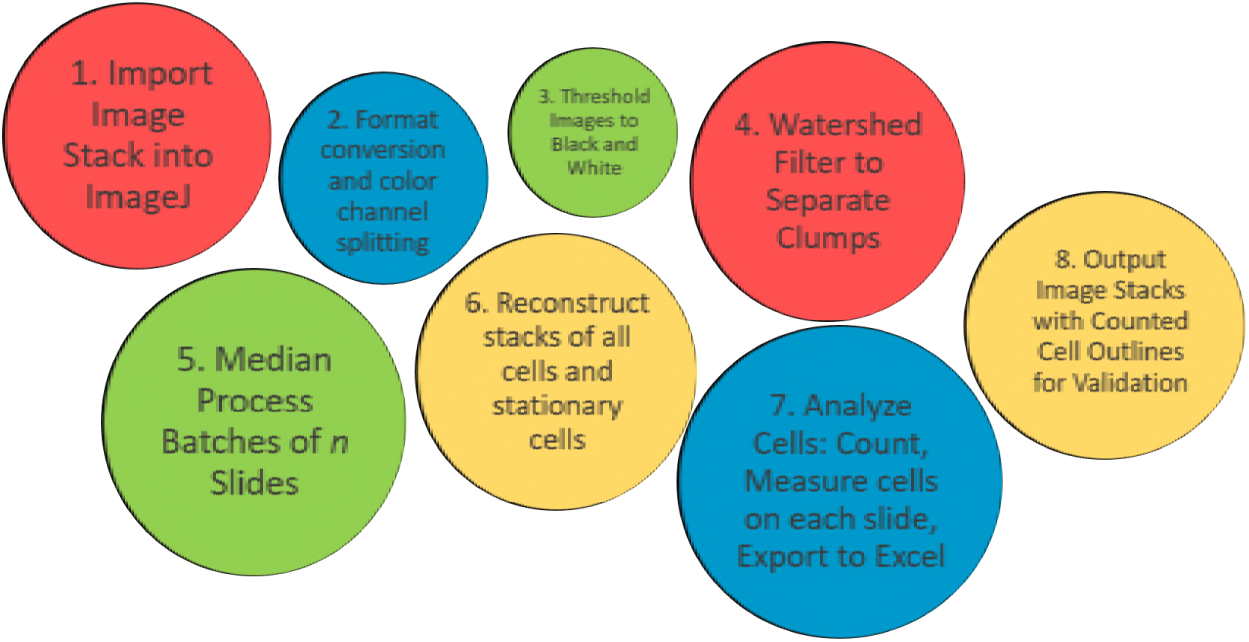
Schematic of macro functions.

In the first step, the user is asked for the pathname for an image stack to be imported and whether the image is single- or multi-channel. Secondly, the color channels (if more than one) are split into distinct image stacks. The user is given the option to choose a thresholding method from standard ImageJ thresholding methods. The program then automatically thresholds all images into black and white. The program then calls another standard ImageJ procedure, watershed, to separate clumps of cells into distinct entities before further processing (11). Next, the program asks the user how many images will be used for the median process, and uses this to run a for loop that calls the process Z Project. In this process, for example, if the user inputs 5 images, then the program will compare images 1-5, assigning each pixel as black if that pixel is black in the majority of the first 5 images, and white if the pixel is white in the first five images. This "median processed" image containing pixels that correspond with the majority of the pixels in the original stack, will be exported into a new stack. The program will then do the same analysis with images 2-6, 3-7, etc. until the end of the stack is reached. At the end, for each image stack, there will be a corresponding median processed stack containing only stationary cells. Next, the program calls the preinstalled ImageJ plugin Analyze Cells. The program asks for 4 parameters: minimum and maximum cell size in pixels, and minimum and maximum cell circularity. Using these parameters, the program calls Analyze Cells to count and measure each cell, produce new image stacks with outlines for each of the cells, and Excel spreadsheets with cell counts and sizes. Finally, the program saves all these files. As a case study, we conducted a DEP separation experiment using a mixture of fibroblasts, stained with calcein green (5*µ*g/mL, Life Technologies), and calcein red (1.7*µ*g/mL, Life Technologies). Further experimental details on chip design, dielectrophoretic theory, or buffer composition, have been discussed at length in our previous publications (7)(6). Prior to this experiment, several voltage, frequency and flow rate combinations were essayed until an optimal combination was selected. Here, at 1.25 *µ*l/min, 20 kHz, and 346 *V_rms_*, the mixed cells are sent through the device. We turned on the voltage, allowing cells to trap, and then turned it off. Occasionally we sped up the flow rate briefly to wash cells through the device. The software OBS (Open Broadcaster Software) was used to record the live view from the Leica X software, and notes were taken when the voltage was turned on/off, cells were washed, etc. This video was exported in .flv format. Using VLC media player scene filter, the video was converted to two stacks of images for further analysis. One stack was sampled at a rate of 6 images per second, for detailed analysis of a short fragment of the video. The same stack was sampled at a rate of 1 image per second for analysis of the whole video.

## RESULTS

Output videos and excel files, the original data set and the macro code are included in the Supplemental Information, along with a user guide. Figures 2, 3 and 4 show an example case of the use of this ImageJ macro. For each of these cases, the gray boxes represent areas with the voltage on, while white is area with the voltage off. The y-axis represents the fraction of cells trapping at each point in time.

**Figure 2:**
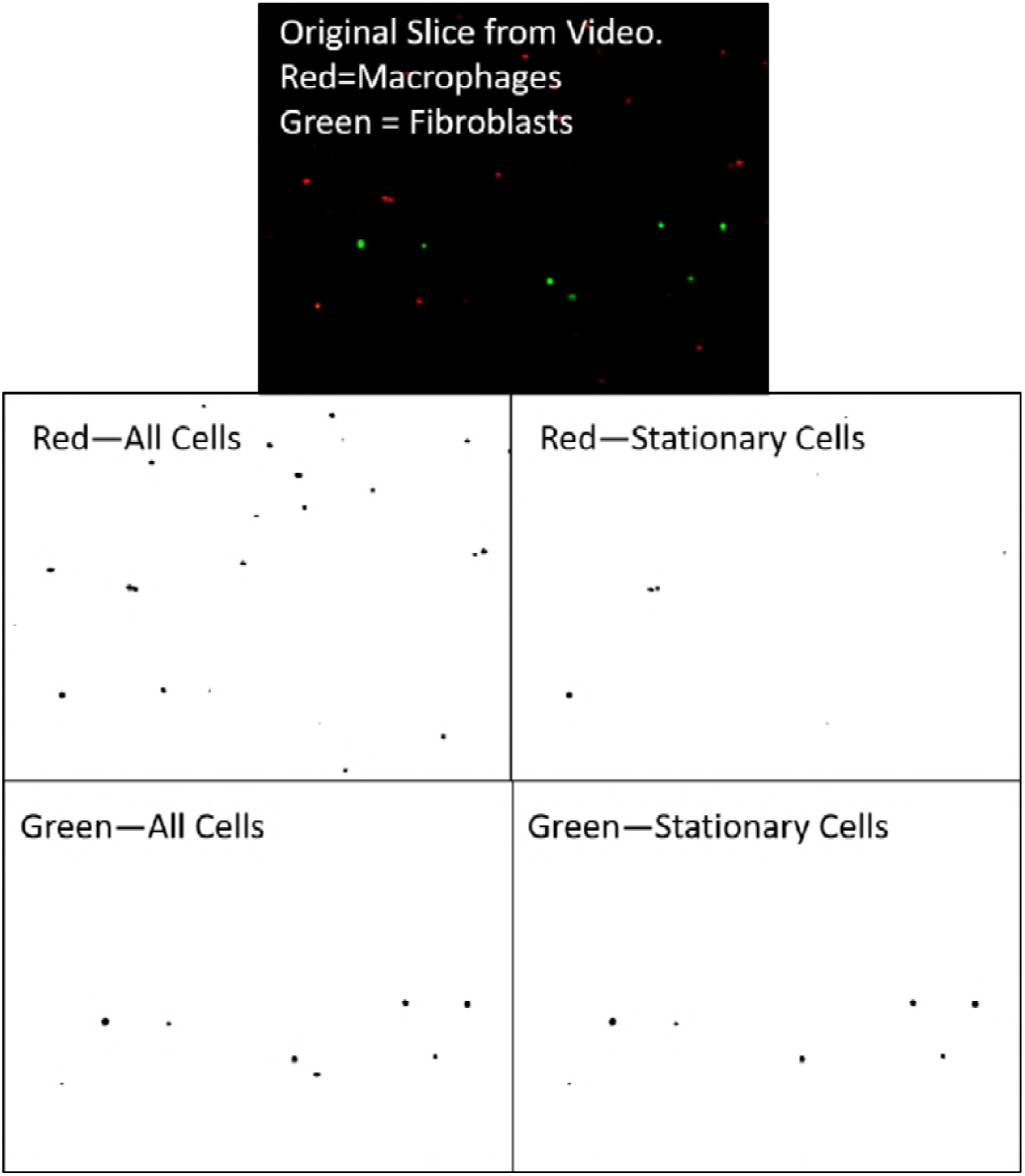
Examples from processed image stack. Videos are available in the supplemental information. The original video slice with fluorescent cells is shown, followed by the split and analyzed images of red and green stationary and total cells for the corresponding median image. Accompanying excel sheets show the cell counts for each slice in the stack.

**Figure 3:**
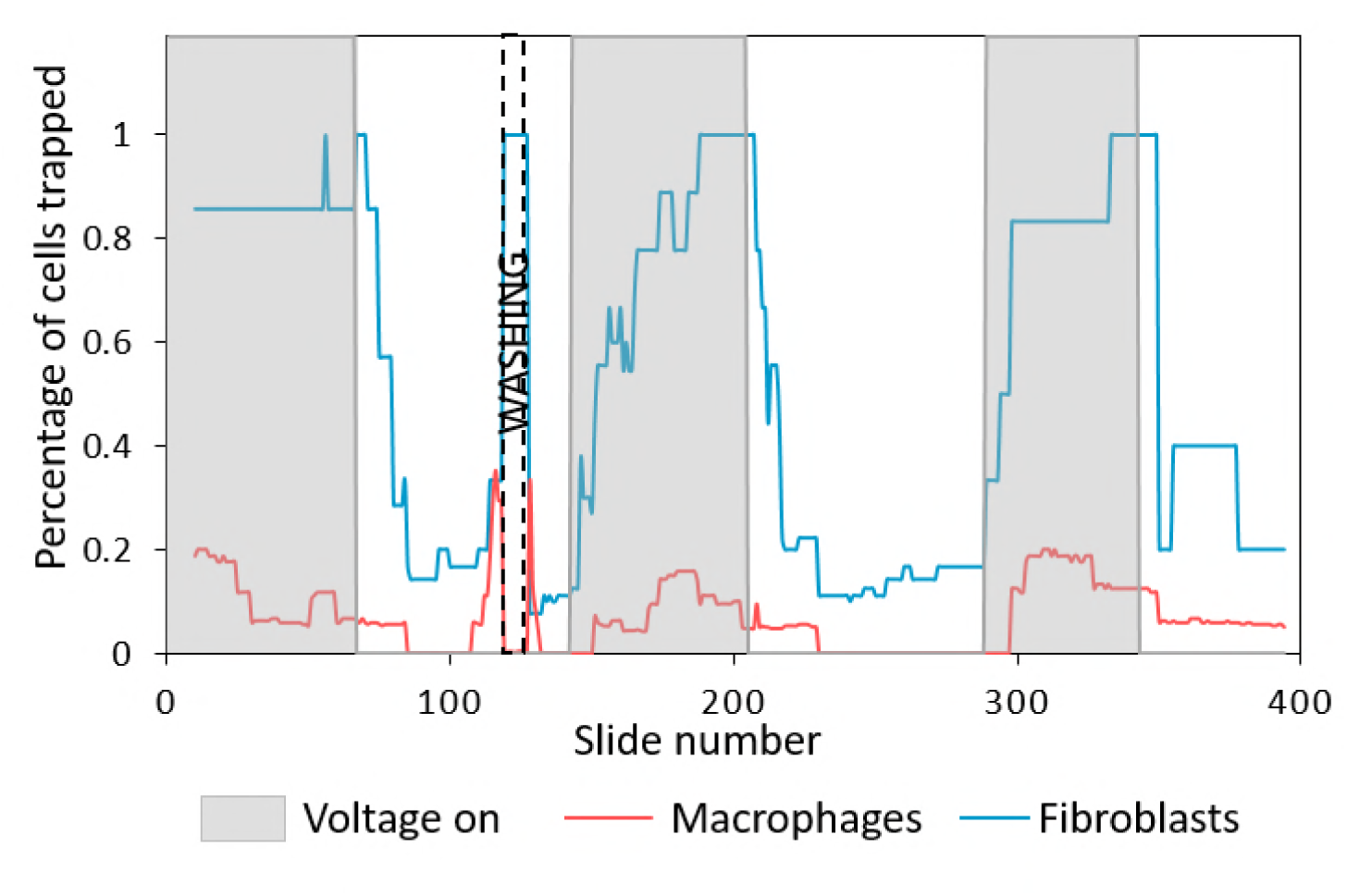
Analyzed data for image stack with 6 frames per second, 21 frames per median process, Otsu filtering, particle size 10-Infinity pixels, particle circularity 0.1-1. Graph shows percent of each cell type trapping at each point in time. As noted, a wash step was performed.

**Figure 4:**
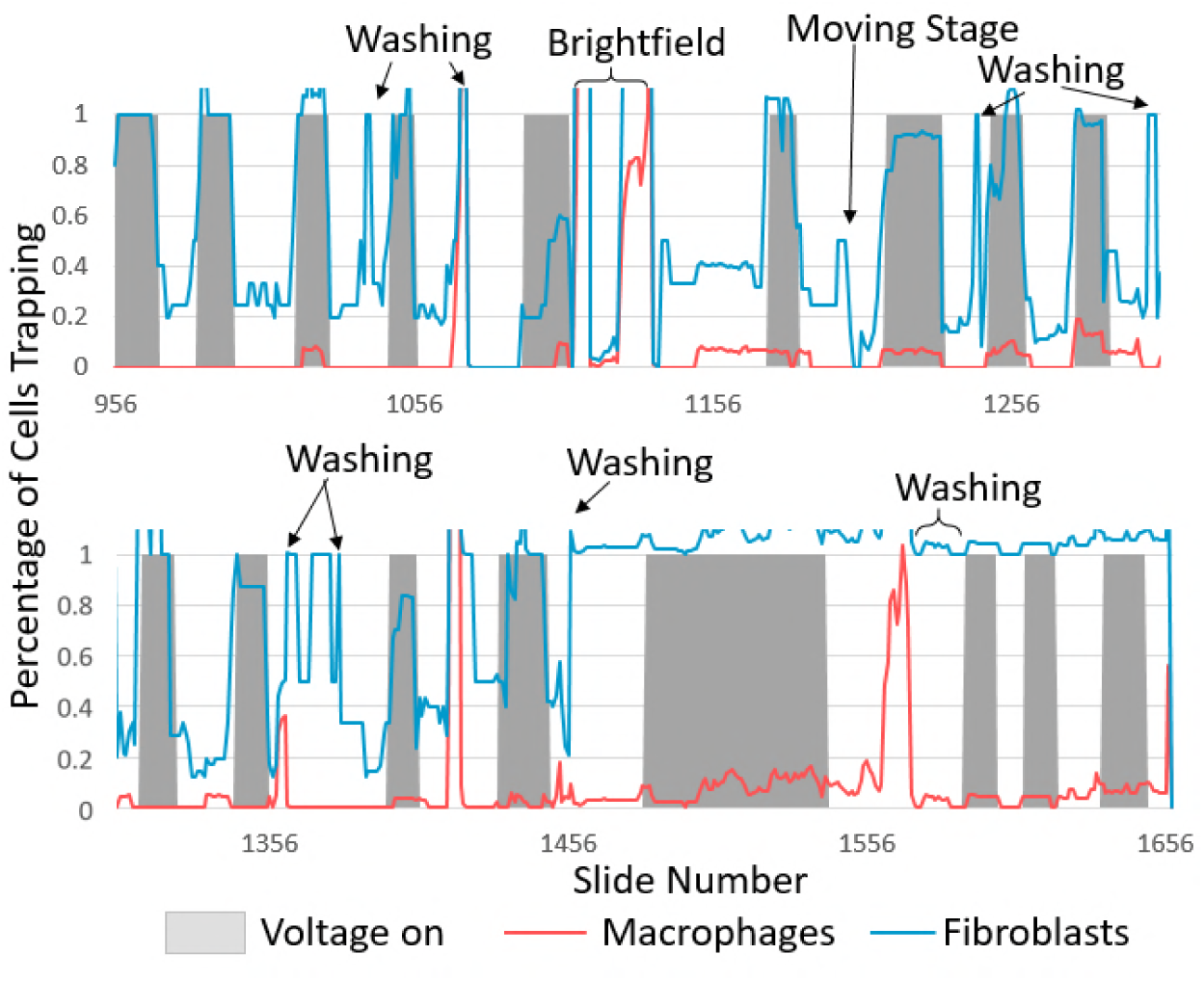
Analyzed data for image stack with 1 frame per second, 7 frames per median process, Otsu filtering, particle size 10-Infinity pixels, particle circularity 0.1-1.(12) Graph shows percent of each cell type trapping at each point in time. Multiple wash steps were performed, brightfield was turned on, and stage was moved to represent different kinds of noise levels.

One drawback to this method is the slight blurring of voltage on-off boundaries by using the median method. When determining whether a cell is moving, the program needs to reference a sequential subset of images. For example, if 15 images are used to determine if the cell is moving, the first image in the median processed (moving vs. stationary) plot is made from taking the majority of the pixels from images 1-15. If, for example, the voltage was turned on between images 3-4, then the first median processed image will show that the cells were trapped. This can be somewhat compensated for by correlating the set with an image in the center of that set, for example, assigning the median image set 1-15 with original image 7, 2-16 with original image 8, etc. One way of analyzing, which was performed in Figures 3 and 4 was to divide the number of cells counted in each median processed image (stationary cells) by the total number of cells in the central image from that median set. This can then be plotted as is shown.

Optimization of the algorithm must be done to ensure reliable results. The standard thresholding methods in ImageJ can be used to test different threshold methods before running the program (13)(14)(15)(16)(17)(18)(19)(20)(21),(22)(12)(23)(24)(25)(26)(27)(28). To optimize the parameters for the median processing and subsequent cell analysis, it may be useful for the researcher to create a file containing a subset of the total number of images for iterative testing, to speed the optimization process. If the number of images chosen is too large, the averaging mechanism may erase some trapped cells. If the number of images chosen is too small, then slow moving particles will show up as trapped. In this case it is also common to see significant levels of noise, where pieces of particles are visible in one frame of the median processing and absent in the next frame.

## DISCUSSION

The development of an ImageJ macro for analyzing videos of moving vs. stationary cells can help to speed and automate the analysis of multiple microfluidic device configurations. It can be very helpful for applications such as ours in the area of dielectrophoresis, where it is necessary to determine percentage of cells trapped by an electric field. (29)

In using this macro, it is necessary to optimize the run parameters before analyzing a full data set, to prevent over or undersampling of data.

In the future, this macro could be developed into a full ImageJ plugin or integrated into video processing software so that cell trapping percentages could be analyzed in real time, allowing researchers to obtain data while running experiments. This could help to speed up the optimization of dielectrophoretic setups where optimizing frequency-voltage pairings is necessary to optimize device functionality.

## CONCLUSION

We have developed an ImageJ installable macro that can be used to count moving vs. stationary objects. As is, it is optimized to be used with moving vs. stationary cells fluorescently labeled cells or particles. However, it is easily modifiable for novel applications, and also comes with user-friendly dialog boxes for easy programming-free input. The code will be listed on the NIH’s macro repository, https://imagej.nih.gov/ij/macros/ (30), and has the DOI 10.5281/zenodo.1204436 at Zenodo, https://zenodo.org/. The code, user guide, and video examples are also found in the supplementary information for this article.

## AUTHOR CONTRIBUTIONS

Temple Douglas designed the macro and tested it. Temple Douglas and Rafael Davalos wrote the article.

## ACKNOWLEDGMENTS

This work was supported by NIH 5R21 CA173092-01. We would also like to acknowledge Virginia Biosciences Health Research Corporation (VBHRC) and NSF IGERT DGE-0966125 MultiSTEPS. This work was supported in part by an unrestricted research grant from CytoRecovery. Davalos has patents in the field of dielectrophoresis.

## SUPPLEMENTARY MATERIAL

An online supplement to this article, with example output videos and a user guide, can be found by visiting BJ Online at http://www.biophysj.org.

